# Unilateral cross-feeding constrains adaptive evolution, even in the producer without direct fitness effects

**DOI:** 10.64898/2026.03.31.715640

**Authors:** Zahraa Al-Tameemi, Thibault Rosazza, Alejandra Rodríguez-Verdugo

## Abstract

Cross-feeding interactions are pervasive in microbial communities and profoundly shape community structure, stability, and function. While previous studies have explored how cross-feeding affects evolvability, this work has predominantly focused on bidirectional mutualistic interactions in engineered auxotrophic systems where both partners reciprocally exchange essential metabolites. However, most metabolic interactions in natural microbial communities are unidirectional, with organisms feeding on the metabolic waste products of other species. Our study addresses this gap by examining how a unidirectional cross-feeding interaction affects the evolutionary dynamics of both the producer (*Acinetobacter johnsonii*) and consumer (*Pseudomonas putida*) over 800 generations of experimental evolution. We found that co-culture constrained adaptive evolution in both species. Co-cultures exhibited lower π_N_/π_S_ ratios (0.75 for *P. putida*; 1.04 for *A. johnsonii*) than monocultures (1.44 and 2.02, respectively) indicating stronger purifying selection against nonsynonymous mutations in the community context. Lineage tracking through whole genome sequencing of populations and clones revealed greater lineage diversity and complexity in monocultures, with more mutations showing significant parallelism across replicate populations. Additionally, *P. putida* evolved increased dependence on its partner; co-culture-evolved *P. putida* grew significantly worse than its ancestor when *A. johnsonii* was removed. These findings demonstrate that ecological interactions fundamentally reshape fitness landscapes and constrain adaptive evolution even when fitness benefits are unidirectional, with implications for understanding microbial community stability and predicting evolutionary dynamics in complex communities.

## Introduction

Microbial populations evolve within communities composed of interacting species. Increasing evidence demonstrates that species interactions are fundamental drivers of evolutionary dynamics and outcomes (Barraclough, 2015; Gorter et al., 2020). By modulating the rate of adaptation (Barber et al., 2022), promoting co-evolutionary feedbacks (Nair et al., 2019) and altering the direction and strength of selection (Adamowicz et al., 2020; Venkataram et al., 2023), species interactions can generate distinct evolutionary trajectories and trade-offs. Elucidating how microbial adaptation occurs within this interactive network is therefore essential for a predictive understanding of evolution in natural microbial communities.

One prevalent form of species interactions in microbial communities is cross-feeding in which one microbe consumes metabolites produced by another microbe. These metabolic exchanges underpin community structure, stability, and biogeochemical function across habitats ranging from the mammalian gut to contaminated soils (Morris et al., 2012; Goldford et al., 2018). A growing body of work has begun to address how cross-feeding interactions influence evolution. Experimental evolution studies with engineered auxotrophic consortia in which both partners reciprocally exchange essential metabolites have demonstrated that obligate mutualistic cross-feeding constrains evolvability: metabolically interdependent organisms are less able to adapt to environmental stress than their free-living counterparts and frequently revert to metabolic autonomy under selection (Pauli et al., 2022; Adamowicz et al., 2018). This constraint has been formalized as the “weakest link hypothesis” which posits that adaptation in mutualistic consortia is limited by the less adaptable partner, because the consortium’s fitness depends on contributions from both members (Melero-Jiménez et al., 2025; Pauli et al., 2022). More recently, Melero-Jiménez et al. (2025) showed that under lethal stress, obligate cross-feeding consortia frequently undergo mutualism breakdown, with one partner regaining autonomy as the primary route to evolutionary rescue (Melero-Jiménez et al., 2025).

Despite these advances, the existing literature is heavily weighted toward bidirectional cross-feeding, typically in engineered systems where both partners exchange essential amino acids or vitamins. However, most metabolic interactions in natural microbial communities are not reciprocal but unidirectional (Freilich et al., 2011). Unidirectional cross-feeding in which one species benefits by consuming the waste products or overflow metabolites from another species, is one of the most prevalent forms of metabolic interactions in microbial communities. These interactions are often commensal, as the producer species does not receive a direct fitness benefit (Zengler and Zaramela, 2018; D’Souza et al., 2018).Whether the evolutionary constraints documented for mutualistic bidirectional cross-feeding extend to commensal unidirectional cross-feeding remains an open question.

Addressing this gap is important for understanding microbial evolution in natural contexts, where unidirectional cross-feeding may be the rule rather than the exception. Such interactions can alter key evolutionary parameters that shape adaptation. For example, in commensal cross-feeding systems the consumer may be constrained by the sporadic release of metabolic byproducts from the producer, resulting in lower population densities. By changing the population size, commensal unidirectional cross-feeding may change the supply of beneficial mutations available to selection (Sniegowski and Gerrish, 2010).

Here, we use a well-characterized commensal cross-feeding system between *Acinetobacter johnsonii* C6 (producer) and *Pseudomonas putida* KT2440 (consumer) to investigate how a unidirectional metabolic interaction shapes evolutionary dynamics over 800 generations of evolution. In this system, *A. johnsonii* oxidizes benzyl alcohol to benzoate as a metabolic byproduct, which is then consumed by *P. putida* as its primary carbon source (Rodríguez-Verdugo et al., 2019; Rodríguez-Verdugo and Ackermann, 2021; Al-Tameemi and Rodríguez-Verdugo, 2024). *P. putida* benefits substantially from co-culture through access to biogenic benzoate, whereas *A. johnsonii* achieves similar population densities regardless of whether its partner is present. Although the effect is subtle, *A. johnsonii* reaches slightly higher densities in monoculture, as it can switch to utilizing benzoate after depleting benzyl alcohol (Rodríguez-Verdugo et al., 2019). In addition to serving as a model for unidirectional cross-feeding, this system reflects natural co-occurrence, as species were co-isolated from hydrocarbon-polluted groundwater in Fredensborg, Denmark (Flyvbjerg et al., 1993; Møller et al., 1996), thereby providing ecological relevance to laboratory investigations of their co-evolutionary dynamics.

We evolved replicate populations of both species in monoculture and co-culture, integrating whole-genome sequencing of populations and individual clones across multiple timepoints with comprehensive phenotypic profiling. This experimental design enabled us to directly compare the mode and tempo of molecular evolution across distinct ecological contexts, track the trajectories of individual mutations, quantify the extent of parallelism as a hallmark of adaptive evolution, and assess how the strength of ecological interactions evolved over the course of the experiment.

We report that co-culture constrains adaptive evolution in both the consumer and the producer species, despite the asymmetric ecological benefits of the interaction. Both species accumulated fewer nonsynonymous mutations, exhibited lower π_N_/π_S_ ratios, and showed reduced parallelism across replicate populations in co-culture compared to monoculture. Lineage tracking of *P. putida* revealed distinct selective sweep dynamics between conditions, with monocultures favoring hard sweeps and co-cultures maintaining greater lineage coexistence. Notably, *P. putida* evolved to become increasingly dependent on *A. johnsonii* over evolutionary time, while *A. johnsonii* shifted toward a slightly antagonistic response to its partner. Additionally, we document the parallel deletion of a 72-kb composite genomic island in *A. johnsonii*, representing convergent genome streamlining independent of the cross-feeding interaction. Collectively, these findings reveal that evolutionary constraints previously documented in obligate mutualisms also emerge in commensal interactions, highlighting the pervasive influence of community context in shaping adaptive trajectories, even when only one partner gains a direct benefit.

## Results

### *A. johnsonii* constrains phenotypic adaptation of *P. putida* over 800 generations

To investigate the effects of species interactions on fitness improvement, we tracked the population density of *A. johnsonii* and *P. putida* in both monoculture and co-culture by counting colony-forming units (CFUs) over 800 generations. *A. johnsonii* maintained a high population density throughout the experiment, with an average density across replicates of 1.2 × 10^8^ CFU mL^-1^ for both monoculture and co-culture conditions (Fig. 1A). Population growth improved over time, with a pronounced increase in density observed between generations 600 and 800. Relative to generation 50, population densities increased significantly by fivefold in monocultures (two-tailed t test, df = 6, *P* value = 0.003) and fourfold in co-cultures (two-tailed t test, df = 6, *P* value = 0.012) by the end of the experiment. Taken together, *A. johnsonii* showed no differences in growth when cultured alone or in the presence of *P. putida*, which is consistent with commensal unidirectional cross-feeding.

**Fig. 1.**
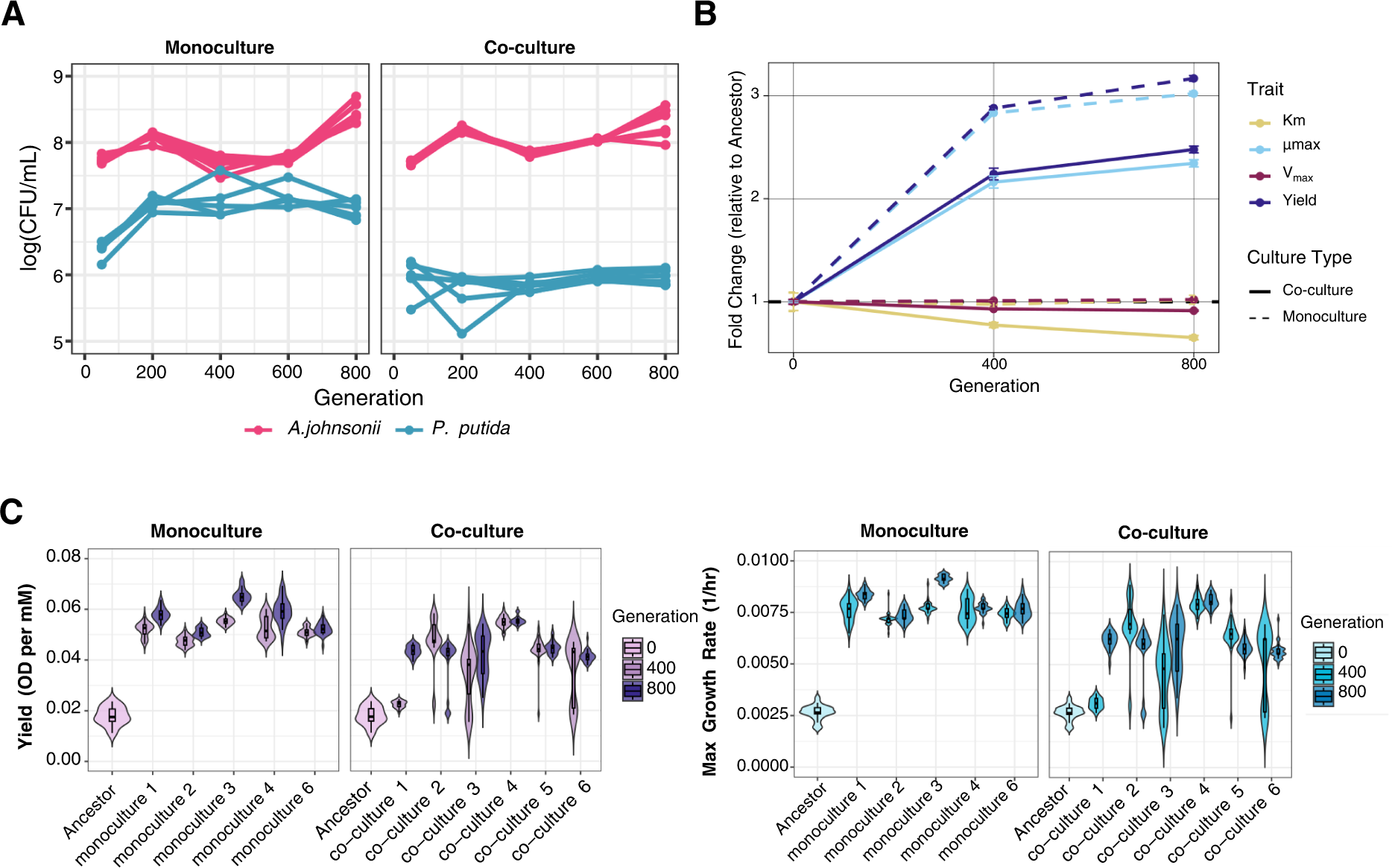
The magnitude of phenotypic improvement in *P. putida* is reduced in co-culture. (A) Population densities (CFU/mL) of *P. putida* (blue) and *A. johnsonii* (pink) in co-culture (left) and monoculture (right) over 800 generations of evolution. (B) Fold change in *P. putida*’s growth parameters −half-saturation constant (*K_m_*), maximum growth rate (*µ_max_*), maximum uptake rate (*V_max_*) and Yield− relative to ancestral values in co-culture and monoculture conditions. At generation 800, monoculture populations of *P. putida* exhibited significantly greater fold changes than co-cultures across all four traits (Welch’s t-test; *Yield*: *P* value = 0.006, *µ_max_*: *P* value = 0.005, *K_m_*: *P* value = 0.012, *V_max_*: *P* value = 0.009; Table S1).” (C) Yield (OD per mM) and maximum growth rate (1/hr) distributions across *P. putida*’s experimental lines at generations 0, 400, and 800. The results for the half-saturation constant (*K_m_*) and maximum uptake rate (*V_max_*) are presented in Figure S2. Pairwise comparisons of these resource use traits across generations are shown in Table S2.

In contrast, we observed striking differences in *P. putida* growth between co-culture and monoculture conditions (Fig.1A). In monoculture, *P. putida* density rapidly increased by an order of magnitude within the first 200 generations of evolution, rising from ∼2.5 × 10^6^ ± 2.9 × 10^5^ CFU mL^-1^ at generation 50 to ∼1.2 × 10^7^ CFU mL^-1^ ± 1.3 × 10^6^ CFU mL^-1^ by generation 200 (Fig.1A). This density was maintained in all independent experimental lines until the end of the experiment at 800 generations. The only exception was monoculture population #5 (PP5) which went extinct after only 50 generations, potentially due to the accumulation of deleterious mutations or stochastic loss during the serial transfers. In co-culture, although all lineages persisted, populations reached densities similar to those at the start of the experiment, averaging ∼9.4×10^5^ ± 9.9 × 10^4^ CFU mL^-1^ at 800 generations (two-tailed t test, *df* = 6, *P* value = 0.736). The lack of significant increase in density over time in co-culture was not due to initial differences in population densities between co-cultures and monocultures (Fig. S1). In the ancestral population, *P. putida* reached a population density of ∼ 1.2×10^6^ ± 4.0×10^5^ CFU mL⁻¹ over four days in co-culture and ∼ 5.6×10^5^ ± 1.9×10^5^ CFU mL⁻¹ in monoculture (two-tailed t test, *df* = 40, *P* value = 0.123; Fig. S1). Thus, although the presence of *A. johnsonii* initially enhanced *P. putida* growth, this advantage did not persist over evolutionary time. Despite the lack of significant growth improvements during the evolution experiment, single clones isolated from co-cultures at generation 800 exhibited a significantly higher yield and maximum growth rate (*µ_max_*) compared to the ancestor when grown on benzoate as the sole carbon source (Fig.1B and C). However, clones from monocultures had a significantly higher yield and maximum growth rate, as well as a significantly lower half-saturation constant (*K_m_*) and maximum uptake rate (*V_max_*), than clones from co-cultures at generation 800 (Fig. 1B, Table S1). Taken together, these findings suggest that interactions with *A. johnsonii* limit growth improvements in *P. putida*.

### Species experience distinct modes of selection when evolved in co-culture compared to monoculture

To get insights into the genetics of adaptation, we sequenced whole populations of *P. putida* and *A. johnsonii* at 800 generations and identified *de novo* mutations (Table S3). When analyzing the mutations fixed in populations or close to fixation (freq >80%) at 800 generations, both species accumulated more nonsynonymous mutations in monoculture than in co-culture (Fig. 2A). This difference was statistically significant for *A. johnsonii* (1.33 vs 0.17 mutations per replicate; Mann-Whitney U test, *P* value = 0.015) but not for *P. putida* (1.8 vs 0.83 mutations per replicate; *P* value = 0.109).

**Fig. 2.**
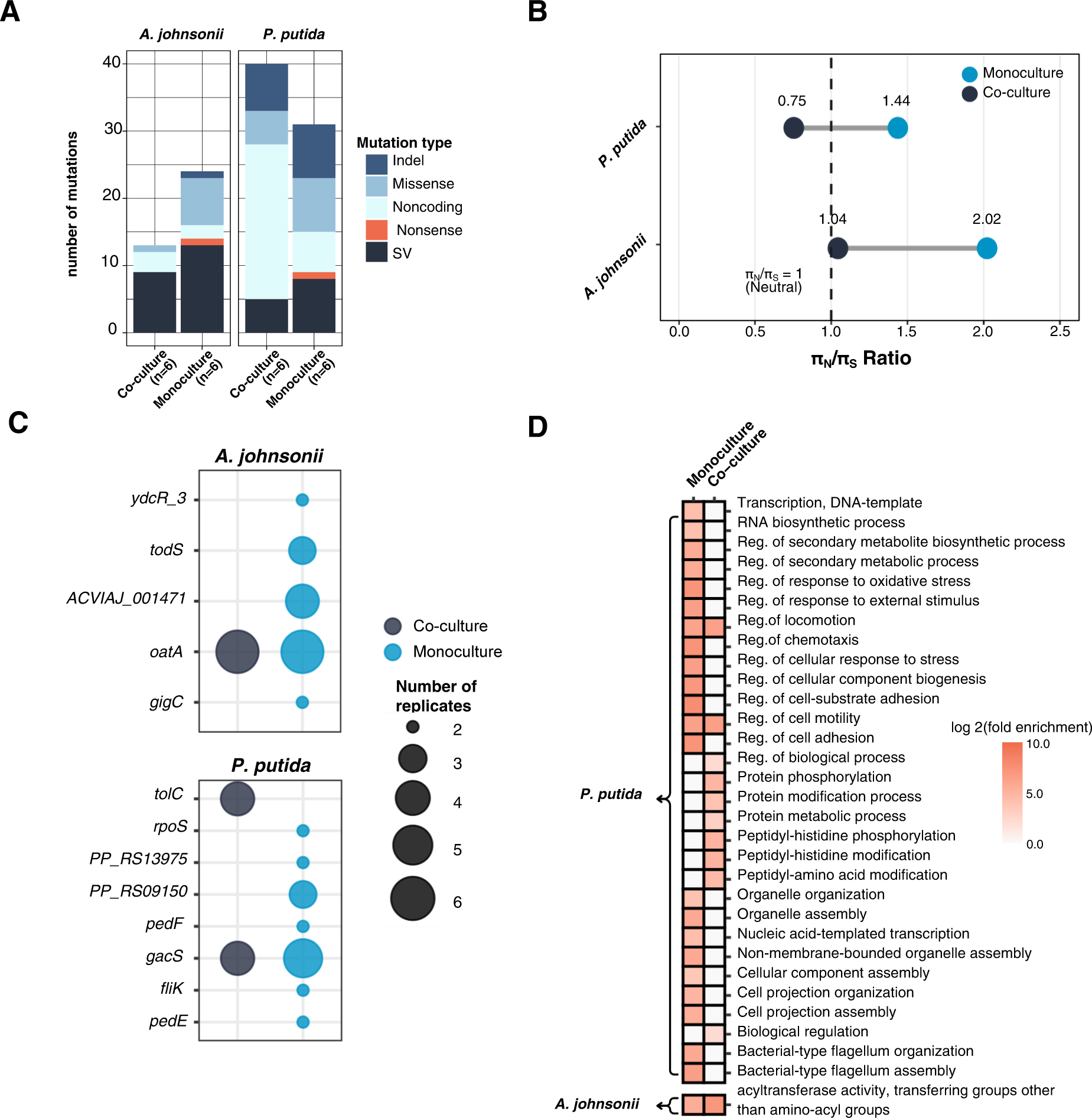
Species exhibit distinct signatures of molecular evolution in monoculture versus co-culture after 800 generations. (A) Mutations count according to types: point mutations in coding regions (with nonsynonymous mutations divided in missense and nonsense mutations), short insertions and deletions (indels), structural variants (SV) and mutations in noncoding regions. (B) π_N_/π_S_ ratios for *P. putida* and *A. johnsonii* in monoculture (blue) and co-culture (dark blue). Dashed line indicates neutral expectation (π_N_/π_S_ = 1). Both species experienced more purifying or neutral selection pressure in co-culture. (C) Mutations show significant parallelism across experimental lines. Circle size indicates number of replicate populations with mutations in each gene. Both species had more mutations significantly enriched by selection (appears across independent lines) in monoculture of both species. (D) GO enrichment analysis of significantly enriched pathways in both conditions. The color saturation indicates the log_2_(fold enrichment) of each pathway.

Next, we investigated whether *P. putida* and *A. johnsonii* experienced different modes of selection in monoculture versus co-culture environments. To do so, we calculated the nucleotide diversity (π) of evolved populations using all detected nonsynonymous (π_N_) and synonymous (π_S_) mutations at generation 800. This analysis revealed striking differences in selection patterns between species and conditions (Fig. 2B, Table S4). In *P. putida* co-cultures, the π_N_/π_S_ ratio was 0.75, indicating purifying selection acting against deleterious nonsynonymous mutations. This contrasted sharply with *P. putida* monocultures, which exhibited a π_N_/π_S_ ratio of 1.44, suggesting relaxed purifying selection or more accumulation of nonsynonymous mutations. This shift from purifying to relaxed selection when *P. putida* was grown in co-culture suggests that the presence of *A. johnsonii* imposed stabilizing selective pressures on *P. putida* populations. This stabilizing selective pressure could explain the lack of significant improvement in growth over time.

*A. johnsonii* experienced similar selective pressures as *P. putida*. In co-culture, it maintained a π_N_/π_S_ ratio near neutral expectations (1.04) and showed the highest π_N_/π_S_ ratio observed across both species and conditions (2.02) in monoculture (Fig. 2B). Taken together, these results show that co-culture environments impose distinct selective pressures on each member of the community even when all other conditions and carbon availability are similar.

### Species show signs of adaptive evolution in both monoculture and co-culture conditions

Despite species experiencing different modes of selection in monocultures and co-cultures, they overall displayed strong signs of adaptive evolution. In particular, we observed pervasive signs of parallel evolution in our experiment. Some genes accumulated mutations across all (or almost all) replicate populations in both monocultures and co-cultures (Table 1). For *A. johnsonii*, the most commonly mutated gene was *oatA* encoding the O-acetyltransferase (OatA) which was mutated in six out of six populations, both monocultures and co-cultures. Mutations in the *oatA* gene appeared early during the evolution experiment and reached fixation by generation 400 in nearly all the replicates (Fig. S3). In *P. putida*, the *gacS* gene, which encodes the sensor protein GacS, accumulated more mutations than expected by chance, indicating strong positive selection (Table 1). Consistent with this observation, *gacS* mutations were fixed early during the evolution experiment (Fig. S3). In monocultures, PP_RS09150, which codes for an acyltransferase family protein, emerged as an additional convergent target (4/5 populations) in *P. putida*. Overall, the strong signature of positive selection on master regulator genes suggests that they play an important role in adaptation to the culture conditions.

**Table 1.**
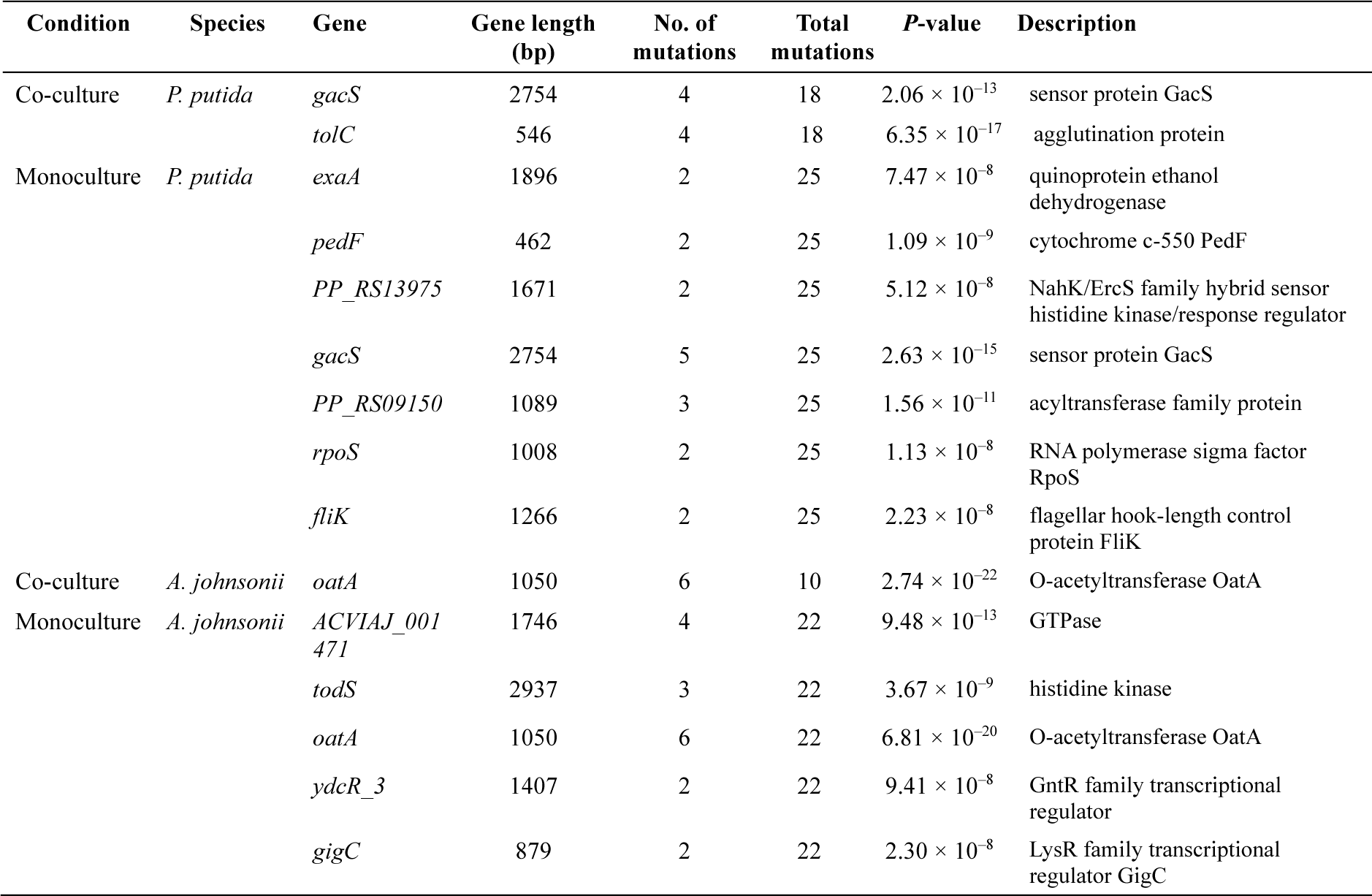
Genes showing significant parallel evolution across independently evolved populations.

| Condition | Species | Gene | Gene length (bp) | No. of mutations | Total mutations | P-value | Description |
| --- | --- | --- | --- | --- | --- | --- | --- |
| Co-culture | *P. putida* | *gacS* | 2754 | 4 | 18 | $2.06 \times 10^{-13}$ | sensor protein GacS |
| | | *tolC* | 546 | 4 | 18 | $6.35 \times 10^{-17}$ | agglutination protein |
| Monoculture | *P. putida* | *exaA* | 1896 | 2 | 25 | $7.47 \times 10^{-8}$ | quinoprotein ethanol dehydrogenase |
| | | *pedF* | 462 | 2 | 25 | $1.09 \times 10^{-9}$ | cytochrome c-550 PedF |
| | | *PP_RS13975* | 1671 | 2 | 25 | $5.12 \times 10^{-8}$ | NahK/ErcS family hybrid sensor histidine kinase/response regulator |
| | | *gacS* | 2754 | 5 | 25 | $2.63 \times 10^{-15}$ | sensor protein GacS |
| | | *PP_RS09150* | 1089 | 3 | 25 | $1.56 \times 10^{-11}$ | acyltransferase family protein |
| | | *rpoS* | 1008 | 2 | 25 | $1.13 \times 10^{-8}$ | RNA polymerase sigma factor RpoS |
| | | *fliK* | 1266 | 2 | 25 | $2.23 \times 10^{-8}$ | flagellar hook-length control protein FliK |
| Co-culture | *A. johnsonii* | *oatA* | 1050 | 6 | 10 | $2.74 \times 10^{-22}$ | O-acetyltransferase OatA |
| Monoculture | *A. johnsonii* | *ACVIAJ_001471* | 1746 | 4 | 22 | $9.48 \times 10^{-13}$ | GTPase |
| | | *todS* | 2937 | 3 | 22 | $3.67 \times 10^{-9}$ | histidine kinase |
| | | *oatA* | 1050 | 6 | 22 | $6.81 \times 10^{-20}$ | O-acetyltransferase OatA |
| | | *ydcR_3* | 1407 | 2 | 22 | $9.41 \times 10^{-8}$ | GntR family transcriptional regulator |
| | | *gigC* | 879 | 2 | 22 | $2.30 \times 10^{-8}$ | LysR family transcriptional regulator GigC |

Two other interesting patterns emerged from the parallelism test. First, monocultures of both species had more parallel mutations across replicates than co-cultures (Fig. 2C). Second, some genes targeted by selection were exclusive to monocultures and co-cultures. For example, in co-culture, *P. putida* accumulated mutations in the *tolC* gene − encoding the outer-membrane protein TolC − in five out of six experimental lines. Importantly, this gene was not mutated in monocultures. Therefore, our results suggest that evolution targets distinct genes in monoculture and co-culture.

Finally, to better understand the functions associated with genes targeted by selection, we conducted a gene ontology enrichment analysis. This analysis revealed shared enrichment of pathways related to motility and biological regulation in *P. putida* across both monoculture and co-culture conditions (Fig. 2D, Table S5). Co-cultures of *P. putida* also showed significant enrichment of protein modification pathways that were not enriched in monocultures. Pathways related to cell projection, organelle assembly, and regulation were additionally enriched in monocultures. Finally, *A. johnsonii* showed significant enrichment of acyltransferase activity under both conditions (Table S5).

### Evolutionary dynamics differ in monocultures and co-cultures over 800 generations

We further investigated the evolutionary dynamics and lineage fate of *P. putida* through whole-genome sequencing of populations and individual clones. We found that lineage competition was rare relative to mutation accumulation within lineages, yet monocultures accumulated more mutations on average than co-cultures (Fig. 3). Monocultures, with an approximately ten-fold larger effective population size and hence greater supply of beneficial mutations, exhibited significantly higher lineage diversity than co-cultures (17.8 ± 6.4 vs. 11.3 ± 2.3 distinct lineages in monoculture and co-culture, respectively; Mann-Whitney U = 25.5, *P* value = 0.032, Cohen’s d = 1.40). Lineage divergence, measured as the maximum number of accumulated mutations within individual lineages, was also greater in monocultures (6.2 ± 1.1 vs. 4.7 ± 1.0 mutations in monoculture and co-culture, respectively; *P* value = 0.022, d = 1.45), consistent with more rapid adaptive walks in larger populations under strong selection.

**Fig. 3.**
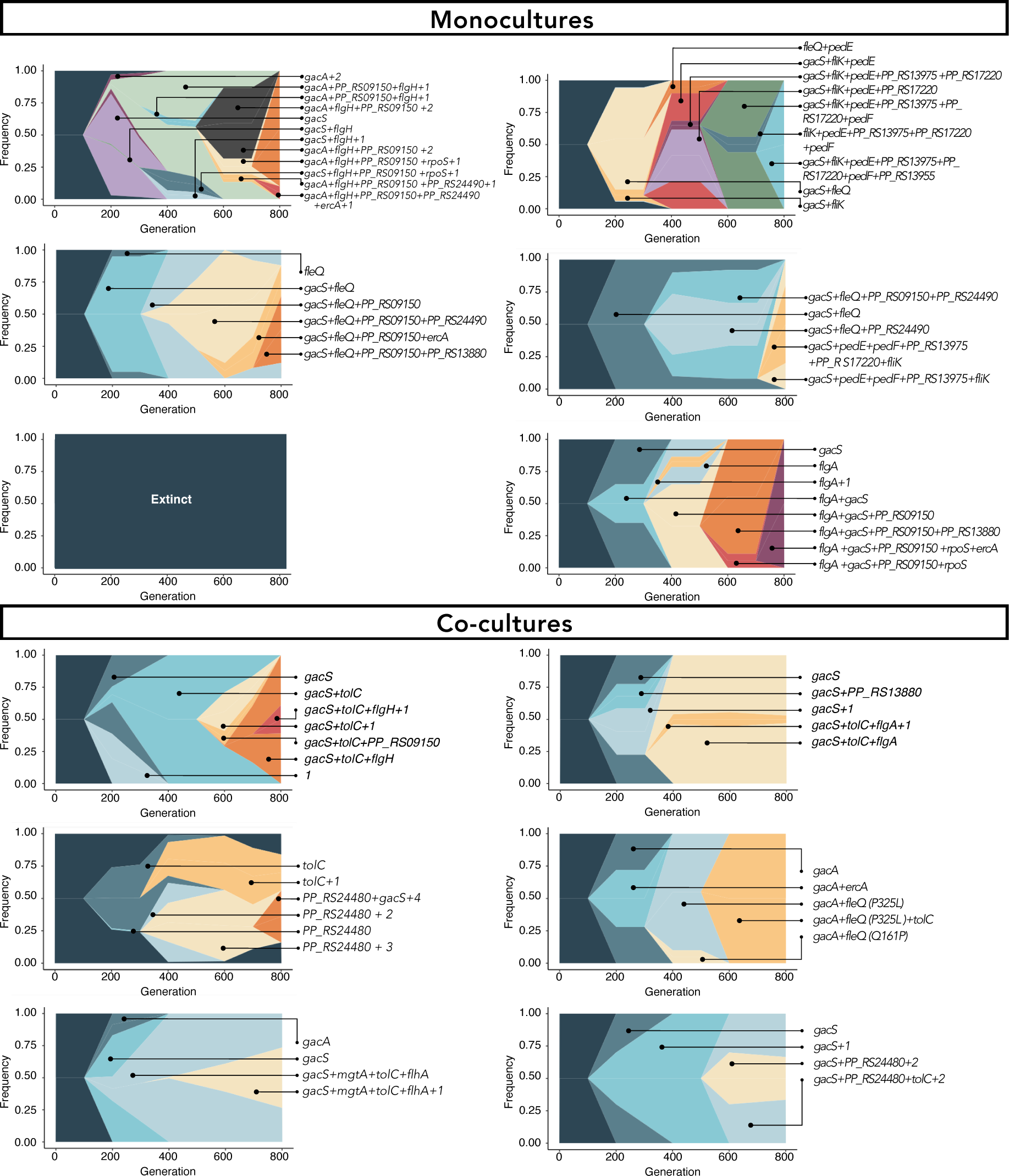
Evolutionary dynamics of *P. putida* are shown using Muller plots. Each color represents a distinct lineage, with nested colors indicating sequential mutation accumulation. The colors do not correspond to specific sets of mutations. Top row: monoculture populations (PP1-6); Bottom row: co-culture populations (Co1-6). Lineage labels indicate the mutational identity (e.g., *GacS*, *tolC*). The mutational labels in the figure include mutations reaching 80% frequency and/or significantly enriched by selection.

*P. putida* experienced distinct selective sweep dynamics between conditions (Fig. 3, Fig. S4). Monocultures exhibited hard sweeps dynamics in three of five populations, with single dominant lineages reaching ≥80% frequency by generation 800 (PP1: 0.90, PP2: 1.00, PP6: 1.00), whereas the remaining populations (PP3, PP4) maintained competing lineages consistent with partial sweeps. In contrast, co-cultures showed a higher proportion of partial sweeps, with four out of six populations (Co1, Co3, Co5, Co6) maintaining multiple coexisting lineages above 10% frequency at the end of the experiment. Notably no single hard sweep was observed in population Co3 (Fig. 3, Fig. S4). These contrasting dynamics were also evident when tracking the fate of mutations reaching over 80% frequency in populations, revealing more hard sweeps in monocultures than in co-cultures for both species (Fig. S4).

Altogether, these results demonstrate that *A. johnsonii* fundamentally reshapes the evolutionary trajectory of *P. putida*, reducing the number of lineages and their complexity while shifting the balance between hard and partial sweeps.

### *A. johnsonii* adapts to the culture conditions through genomic streamlining

In addition to point mutations, other types of mutations arose during our evolution experiment. Notably, large deletions were recurrently observed during *A. johnsonii’*s evolution. We identified a 72-kb composite genomic island at the tmRNA ssrA locus that was independently deleted in 50% of evolved *A. johnsonii* populations. Three distinct genomic islands were identified within this region (Fig. 4, Table S6). Island 1 (coordinates 856,233-861,527; 5.3 kb) contains phage-related genes, including a YqaJ viral recombinase. Island 2 (coordinates 889,088-900,395; 11.3 kb) contains an OmpA family protein and a large BapA/Bap family biofilm-associated protein. Finally, Island 3 (coordinates 908,174-921,589; 13.4 kb) contains metabolic genes including oxidoreductases, dehydrogenases, and an MFS transporter. This deletion was fixed in three monocultures and three co-cultures at generation 400. Excision occurred via site-specific recombination between 16 bp direct repeats (CGCCACCTCCACCAAA) flanking the entire 72-kb region, consistent with prophage-like excision mechanisms (Grindley et al., 2006).

**Fig. 4.**
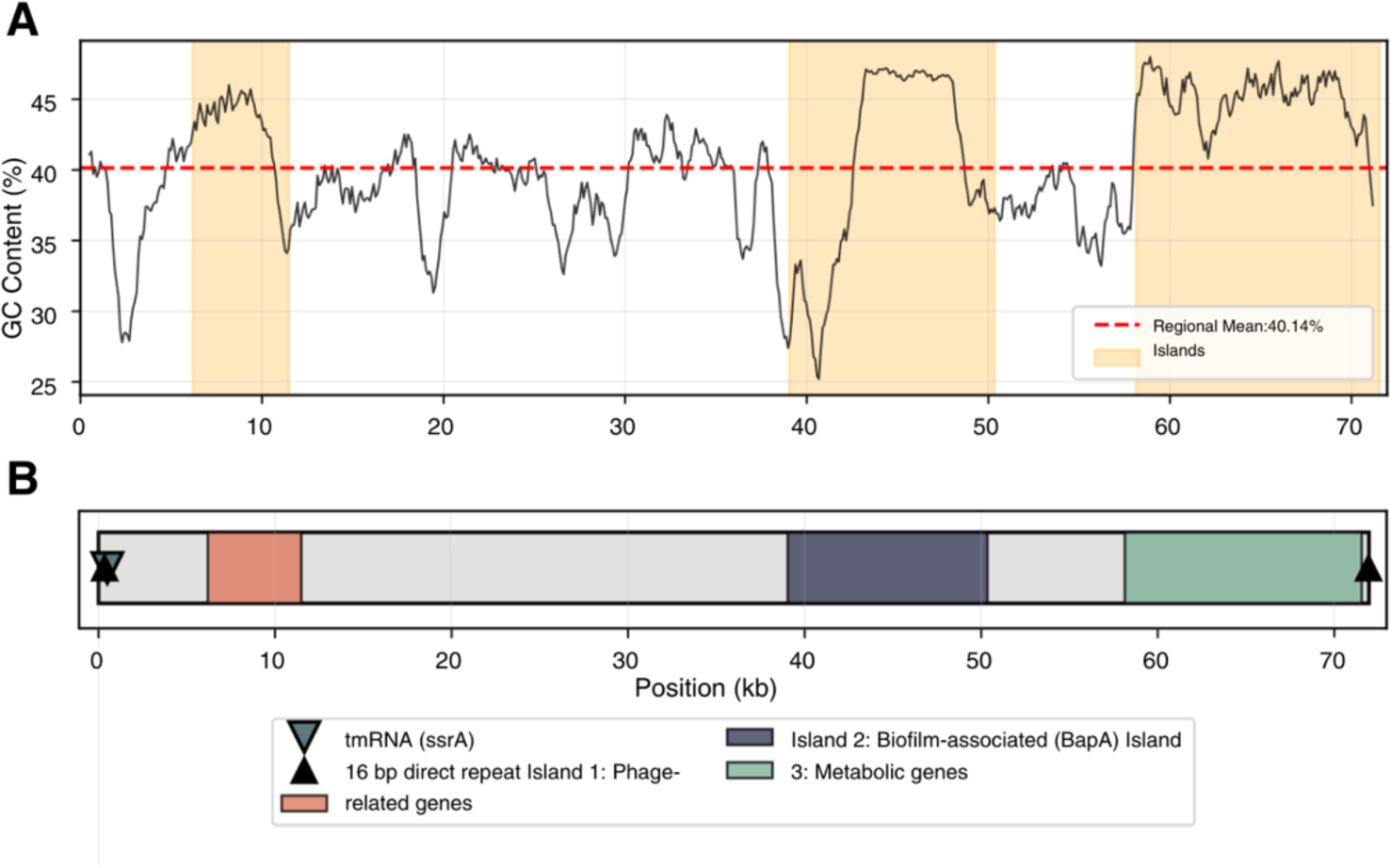
Genomic island structure in *A. johnsonii* C6. (A) GC content variation across the 72-kb deleted region, with genomic islands highlighted. (B) Map of the composite genomic island showing Island 1 (phage-related), Island 2 (biofilm-associated/BapA), and Island 3 (metabolic genes). Direct repeats (16 bp) flanking the region are indicated.

Compositional analysis revealed that the combined genomic islands exhibited significantly elevated GC content (43.0%) compared to the overall extracted region (40.1%) and flanking sequences (χ² = 80.3, p < 0.0001), consistent with horizontal acquisition from a donor organism with higher genomic GC content than *A. johnsonii* C6. Island 3 showed the strongest signature of foreign origin with 45.2% GC content. GC skew analysis showed relatively consistent values across the region (mean = 0.101 ± 0.042), suggesting the island had undergone amelioration toward host codon usage patterns over evolutionary time.

In summary, the independent deletion of this 72-kb composite genomic island in 50% of evolved populations from both monoculture and co-culture conditions suggests strong selection for genome streamlining under laboratory conditions, where the metabolic burden of maintaining biofilm-associated genes and other island-encoded functions likely outweighed any potential fitness benefits.

### Species interactions strengthen over evolutionary time

The ancestral consortium was previously shown to exhibit a commensal interaction when grown on benzyl alcohol as the sole carbon source, with *P. putida* growing ten times better with *A. johnsonii* than alone (Rodríguez-Verdugo et al., 2019; Rodríguez-Verdugo and Ackermann, 2021). To determine whether this interaction changed over 800 generations, we revived co-cultures from generations 6, 400 and 800 and selectively removed either *P. putida* or *A. johnsonii*. This allowed us to compare the growth of monocultures and co-cultures over four passages using benzyl alcohol as the sole carbon source (see Materials and Methods).

We observed that *P. putida* grew better in co-culture than in monoculture across all generations, despite an overall decline in population densities over the 4-days experiment (Fig. S5). Importantly, *P. putida* from 800-generations co-cultures grew better with *A. johnsonii* than *P. putida* from the 6- or 400-generations co-cultures (Fig. S5, Table S7). This growth improvement amplified the difference between co-cultures and monocultures, indicative of a stronger partner’s effect on growth and therefore a stronger interaction (Fig. 5). *P. putida* started with a co-culture advantage of +2.17 AUC units at 6 generations, which nearly doubled to +4.06 AUC units by 800 generations, representing a significant increase in co-culture advantage over evolutionary time (slope = +0.0024 AUC units/generation, *P* value = 2.27×10⁻⁶; Table S8).

**Fig. 5.**
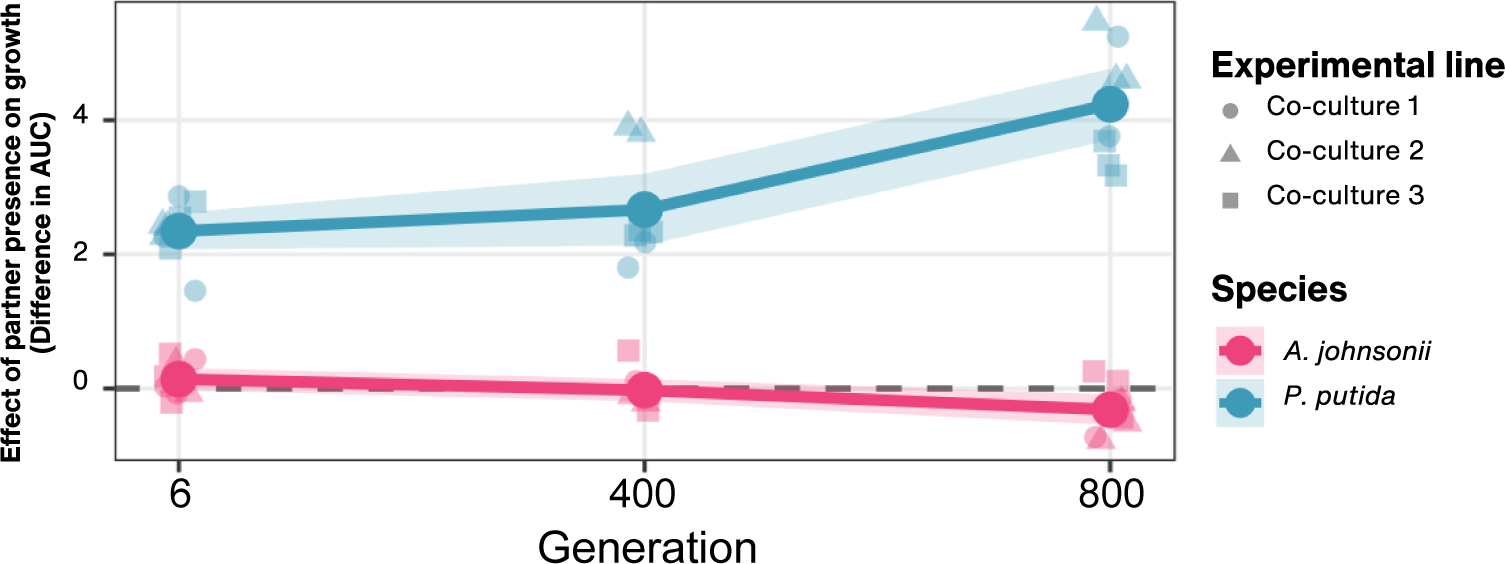
Species interactions intensify over time, with *P. putida* evolving increased benefits from *A. johnsonii,* while *A. johnsonii* incurring fitness costs from the interaction. The effect of the partner species on the growth of the other species was measured as the difference in AUC (area under the curve) between co-culture and monoculture conditions (Table S7).

Interestingly, *A. johnsonii* exhibited an opposite trend, with a significant decrease in co-culture advantage over evolutionary time (slope = -0.00057 AUC units/generation, *P* value = 0.0021; Table S8). *A. johnsonii* started with no significant co-culture advantage at 6 generations (+0.14 AUC units, one sample t-test, *P* value = 0.141), becoming a significant co-culture disadvantage by 800 generations (-0.32 AUC units, one sample t-test, *P* value = 0.027). In conclusion, *P. putida* evolved to grow increasingly better with *A. johnsonii,* while *A. johnsonii*’s growth became slightly inhibited by *P. putida*.

## Discussion

Cross-feeding interactions are pervasive in microbial communities and profoundly shape community structure, stability, and function (D’Souza et al., 2018). While previous studies have explored how cross-feeding affects evolvability, this work has predominantly focused on bidirectional, mutualistic interactions—typically engineered auxotrophic systems where both partners reciprocally exchange essential metabolites (Pauli et al., 2022; Adamowicz et al., 2018; Preussger et al., 2020). However, most metabolic interactions in natural microbial communities are unidirectional, with organisms feeding on the metabolic waste products or overflow metabolites of other species in commensal arrangements (Zengler and Zaramela, 2018; D’Souza et al., 2018). Furthermore, even when interactions appear ecologically neutral for one partner, the evolutionary consequences of co-occurrence may be substantial if species have adapted to each other’s presence. Our study addresses this gap by examining how a unidirectional, commensal cross-feeding interaction affects the evolutionary dynamics of both the producer (*A. johnsonii*) and consumer (*P. putida*) species over 800 generations of experimental evolution.

### Adaptive evolution is constrained in co-culture at both phenotypic and genotypic levels

A central finding of our study is that species experienced lower rates of adaptive evolution in co-culture conditions. Several lines of evidence support this trend. First, *P. putida* exhibits markedly reduced growth improvements in co-culture compared to monoculture. Second, species displayed an elevated π_N_/π_S_ ratios in monocultures compared to co-cultures. Third, both species had more hard sweeps and more parallel mutations enriched by selection across replicates than co-cultures. Fourth, *P. putida* showed greater lineage diversity and complexity in monocultures. Collectively, these results suggest that adaptive evolution is constrained for both *A. johnsonii* and *P. putida*. Adaptive constraints driven by interspecies interactions have been reported in multiple studies, with differences in population size commonly proposed as an explanation for this pattern (Barber et al., 2022; Adamowicz et al., 2020; Castledine et al., 2020; Castledine et al., 2025).

In our system, the constrained adaptive evolution observed in co-cultures could also be explained by differences in population size. *P. putida* in monoculture achieved population densities approximately one order of magnitude higher than in co-culture, which increases the supply of beneficial mutations and accelerates adaptation (Sniegowski and Gerrish, 2010). However, the lower population size in co-culture alone cannot fully explain the reduced π_N_/π_S_ ratio, as this same pattern held for *A. johnsonii* despite achieving similar densities in both conditions. This observation reinforces our hypothesis that ecological context, rather than population size alone, drives the evolutionary constraints observed.

Ecological interactions may constrain adaptive evolution by imposing stabilizing selection pressures. Consistent with this idea is the “weakest link hypothesis” which provides a theoretical framework for understanding reduced evolvability in interacting species (Melero-Jiménez et al., 2025; Pauli et al., 2022; Adamowicz et al., 2018; Adamowicz et al., 2020). This hypothesis posits that adaptation in mutualisms is constrained because the consortium’s fitness is limited by the less adaptable partner. While a single strain can adapt with a single beneficial mutation, a mutualistic consortium requires both partners to adapt or for one to compensate (Melero-Jiménez et al., 2025). Although originally formulated for obligate mutualisms, our results suggest that an analogous constraint may operate even in commensal interactions. The presence of an interacting partner, regardless of whether the interaction confers direct fitness benefits, may stabilize the adaptive landscape and reduce the selective advantage of novel mutations.

Perhaps the most surprising finding of our study is that the evolutionary constrain imposed by co-culture extends to *A. johnsonii*, the producer species that receives no direct fitness benefit from the interaction. This finding challenges the intuitive expectation that only species directly affected by an interaction should experience altered evolutionary dynamics. We propose that even in the absence of strong direct fitness effects, the presence of an interacting species fundamentally alters the selective environment. *A. johnsonii* in co-culture experience a different ecological context: its waste product (benzoate) is rapidly consumed by *P. putida*, potentially altering local metabolite concentrations, redox conditions, or other aspects of the chemical environment. Our results agree with other studies showing that, even when species interactions do not produce major effects on growth, they can still drive major changes in genomic evolution (Putnam et al., 2025).

### Targets of selections differed between species and conditions

Despite reduced overall adaptive evolution in *P. putida* co-cultures, certain mutations showed remarkable convergent evolution across conditions. Mutations in the GacS/GacA two-component regulatory system appeared in nearly all *P. putida* populations of both conditions. The GacS/GacA system is a master regulator controlling secondary metabolism, biofilm formation, motility, and stress responses in *Pseudomonas* species (Heeb and Haas, 2001; Lapouge et al., 2008). The universal targeting of this regulatory system likely reflects adaptation to laboratory culture conditions rather than the specific selective pressures of cross-feeding. Similar convergent mutations in GacS/GacA have been observed in other experimental evolution studies with *Pseudomonas*, often associated with shifts between planktonic and biofilm lifestyles or alterations in secondary metabolite production (Yan et al., 2018; Brencic and Lory, 2009).

Interestingly, while regulatory genes were targeted in both conditions, distinct sets of genes were uniquely targeted in monocultures and co-cultures. Notably, mutations in *tolC* were only observed in co-culture conditions. TolC is part of multi-component efflux pumps in the outer membrane and has been shown to facilitate the export toxic compounds from the cell in other bacterial species (Koronakis et al., 2000; Koronakis, 2003; Langevin and Dunlop, 2018). It is unclear what the function of TolC is under our experimental conditions, but given the observed parallelism in co-cultures, we hypothesize that these mutations represent adaptations to the interaction with *A. johnsonii*. In sum, this differential targeting suggests that selective pressures differ sufficiently between conditions to favor distinct adaptive paths, even when master regulators are universally mutated.

In the case of *A. johnsonii*, populations evolved in monoculture and co-culture shared deletions of a 72-kb composite genomic island in 50% of populations. This represents a striking example of convergent genome streamlining, which was not observed in *P. putida* lines, where parallel mutations were mainly non-synonymous mutations and indels. This deletion, occurring via site-specific recombination between flanking direct repeats, removed genes associated with phage functions, biofilm formation, and various metabolic processes. The elevated GC content of the deleted region (43% vs. 40% for flanking regions) is consistent with horizontal acquisition from a donor with higher genomic GC content, and the parallel loss across multiple independent populations suggests strong selection against maintaining these foreign genetic elements under laboratory conditions.

Genome streamlining through deletion of horizontally acquired elements is a common feature of laboratory evolution, reflecting the removal of conditionally beneficial genes that impose metabolic costs in the stable environment of serial batch culture (Giovannoni et al., 2014; Morris et al., 2012). The fact that this deletion occurred with equal frequency in monocultures and co-cultures suggests it represents an adaptation to general laboratory conditions rather than to the presence or absence of the cross-feeding partner.

### Species interactions slowly changed during the evolution experiment

Species interactions are not static and have been shown to evolve over time in both their nature and their strength (Hansen et al., 2007; Andrade-Domínguez et al., 2014; Fiegna et al., 2015). While the nature of the interaction remained commensal throughout the evolution experiment, its strength intensified (Fig. 5). *P. putida* evolved increased benefits from *A. johnsonii* in co-culture, whereas *A. johnsonii* evolved increased costs in the presence of *P. putida*. This finding aligns with previous observations indicating that *P. putida* exerts a slight but significant negative effect on *A. johnsonii,* consistent with an exploitative interaction (Rodríguez-Verdugo et al., 2019). Interestingly, we observed that *P. putida* evolved a higher uptake at low nutrient concentrations, reflected by a shift toward a lower *K_m_* (Fig. 1B; Fink et al., 2023), with this trait changing significantly only under co-culture conditions. This may explain how *P. putida* becomes more efficient at extracting benzoate from *A. johnsonii*, thereby imposing a greater cost on *A. johnsonii*, which lacks access to this additional benzoate. Consistent with this trend, a previous study found that interactions within this consortium rapidly shifted from commensalism to exploitation when species were grown in biofilms (Hansen et al., 2007). Remarkably, this change in interaction evolved in only ten days, compared to our experiment, in which the trend emerged only after four months. These differences may reflect the influence of spatial structure, suggesting that interactions evolve more rapidly when species are in close proximity, as in biofilms, compared to liquid culture conditions with constant shaking. It is also important to consider that the quantification of interaction strengths was performed using populations revived from frozen glycerol stocks, and the freeze-thaw process may affect population dynamics and species ratios in ways that do not perfectly recapitulate the conditions during the evolution experiment. While we attempted to minimize these effects through acclimation periods, they represent unavoidable technical constraints of working with mixed microbial communities.

In conclusion, the patterns of constrained adaptive evolution in co-cultures of both species suggest that community context fundamentally reshapes the fitness landscape. When evolving within a community, both species may experience a more constrained set of viable evolutionary paths, regardless of the direct fitness consequences of the interaction. This has important implications for understanding microbial evolution in natural environments, where species rarely evolve in isolation.

## Materials and Methods

### Growth conditions

All strains were grown in 10 mL of FAB minimal medium, composed of 1 mM MgCl₂, 0.1 mM CaCl₂, 0.003 mM FeCl₃, 15 mM (NH₄) ₂SO₄, 33 mM Na₂HPO₄, 22 mM KH₂PO₄, and 50 mM NaCl, supplemented with 0.6 mM benzyl alcohol. In *P. putida* monocultures, 0.06 mM benzoate was additionally supplemented to match the concentration naturally produced by A. johnsonii (Rodríguez-Verdugo et al., 2019). The medium was sterilized using a 0.2-µm filter and stored in glassware treated to remove assimilable organic carbon (AOC), following the protocol described by Rodríguez-Verdugo et al. 2019 (Rodríguez-Verdugo et al., 2019). All cultures were incubated at 26°C with constant shaking at 150 rotations per minute (rpm).

### Evolution experiment

*Pseudomonas putida* KT2440 and *Acinetobacter johnsonii* C6 were evolved either separately, in monocultures, or together in co-cultures for approximately 800 generations. Co-cultures and monocultures of *A. johnsonii* were supplemented with benzyl alcohol (0.6 mM). *P. putida* monocultures were additionally supplemented with 0.06 mM of benzoate, to control for the amount of benzoate produced by *A. johnsonii* in co-cultures (Rodríguez-Verdugo et al., 2019). For each treatment (monoculture and co-culture), six replicates were evolved in parallel in FAB minimal medium supplemented with the appropriate carbon sources. Cultures were propagated by daily transferring 0.1 mL of culture into 9.9 mL of fresh medium for 5 months (∼800 generations).

The evolution experiment was started as follows. *P. putida* and *A. johnsonii* were streaked onto LB agar plates supplemented with gentamicin (10 µg/mL) and streptomycin (64 µg/mL), respectively, and incubated at 30°C. A single colony of each species was randomly selected and inoculated into 10 mL of FAB medium with the aforementioned carbon source concentrations. Cultures were incubated at 26°C with constant shaking at 150 rpm for 24 hours. All experimental lines were started from these two single colony cultures. Monocultures were launched by inoculating 100 µL of cultures of either *P. putida* or *A. johnsonii* into 9.9 mL of FAB medium with the previously mentioned concentration of carbon sources. Co-cultures were initiated by mixing 50 µL from each species into 9.9 mL of FAB medium. We confirmed equal starting ratios by plating all cultures immediately following inoculation to insure a 1:1 starting ratio of *P. putida* to *A. johnsonii*. Glycerol stocks of the initial single-colony cultures were preserved as time point zero for subsequent analyses.

To capture growth dynamics during the evolution experiment, *P. putida* and *A. johnsonii* densities were estimated by plating fresh cultures onto selective LB agar with gentamicin or streptomycin every 200 generations. We additionally plated all populations from the evolution experiment weekly to monitor possible contamination or cross-contamination. Glycerol stocks of all populations were saved every 200 generations for subsequent phenotypic and genomic analyses.

### Phenotypic characterization of evolved populations of *P. putida*

To characterize changes in growth parameters of *P. putida* in monoculture and co-culture, we measured the growth curves of 30 single clones of the ancestral *P. putida* population at generation 0 and each experimental line at generations 400, and 800 as in Al-Tameemi & Rodríguez-Verdugo (2024). Briefly, bacteria were grown in FAB medium supplemented with 0.6 mM benzoate for 24 h and were then plated on Lysogeny Broth (LB) agar and incubated for another 24 h at 26°C. We randomly selected 30 colonies and resuspended them into wells filled with 200 µL 1% MgSO4 in a 96-well plate. We then transferred 2 µL of the diluted cells into another 96-well plate containing 198 µL of FAB supplemented with 0.6 mM benzoate. The plate was sealed with vacuum grease to reduce evaporation. The plate was incubated in a photospectrometer plate reader (Epoch2, Agilent) at 30°C and continuous linear shaking. Optical density (OD) measurements were taken every 10 min for 24 h.

From these growth curves, we estimated four quantitative traits involved in resource use: the maximum growth rate in exponential phase (*µ_max_*), the yield (*Y*) defined as the amount of biomass produced per unit of resource, the maximum uptake rate (*V_max_*) defined as the maximum rate of resource consumption per hour and the half-saturation constant (*K_m_*) defined as the resource concentration supporting half-maximum uptake rate as in Rodríguez-Verdugo et al. 2019. Parameters were fitted to the growth curve of each colony using a custom python script (version 3.14).

To test for significant changes in traits across generations, we used linear models with generation (0, 400, 800) as a fixed effect, analyzed separately for each evolved population. Pairwise comparisons between timepoints (generation 0 vs. 400, generation 0 vs. 800, and generation 400 vs. 800) were performed using estimated marginal means with no adjustment for multiple comparisons. Fold changes were calculated relative to ancestral (generation 0) mean values across all populations within each culture type. All statistical analyses were performed in R (version 4.4.2) using the *emmeans* package.

### Library preparation and genome resequencing

Whole populations were sequenced by reviving frozen glycerol stocks from all experimental lines of *P. putida* and *A. johnsonii* at generations 0, 200, 400, 600, and 800. Cultures were launched by transferring 100 µL of glycerol stock into fresh FAB media containing the same carbon sources and conditions under which the populations evolved. The size of the glycerol stock inoculum avoids a population bottleneck and ensures that the revived samples are representative of the genetic diversity present at each generational timepoint. Cultures were grown 24 h at 26°C with constant shaking (150 pm). Genomic DNA was extracted from 1 mL of culture using the Wizard DNA purification kit (Promega).

We additionally isolate eight clones of *P. putida* from monocultures and co-cultures at each of the timepoints to track the evolutionary dynamics of *P. putida* in both conditions. To do so frozen glycerol stocks we revived (see above) and plated on LB agar and incubated for 24 h at 26°C. We randomly selected eight clones off the plate and resuspended them in 3 mL of LB broth and grown overnight at 26°C with constant shaking (150 rpm). Clones from the two ancestors *A. johnsonii* and *P. putida* were directly revived from glycerol stocks and grown overnight (26°C, 150 rpm) in 3 mL LB broth. We isolated and purified genomic DNA from 1 mL of overnight cultures using the PureLink Pro 96 Genomic DNA Purification Kit.

We prepared libraries for whole genome sequencing by using a low volume protocol with the Illumina DNA prep kit (Weihe and Avelar-Barragan, 2021). The libraries were then sent to the UCI Genomics High Throughput Facility for short-read sequencing using the Illumina NovaSeq X Plus platform targeting fragment size of 2×150 bp and 100x coverage.

### Sequence quality control and Variant calling

To ensure sequence quality and absence of adapter contamination, we used *fastp* (Chen et al., 2018) to trim the adapters and low-quality reads with a score lower than 30. The removal of low-quality reads and adapters was confirmed using *multiqc* (Ewels et al., 2016). Whole population sequences from the evolution experiment were then aligned to *P. putida* KT2440 (reference NC_002947) and *A. johnsonii C6* (reference CM142178.1) using the computational pipeline *breseq* (version 0.39.0) and utility program GDtools (Deatherage and Barrick, 2014). Sequence alignment and variant calling was done in consensus mode for clones and polymorphism mode for populations. The ∼100x coverage enabled us to confidently call mutations above 5% frequency. *De novo* mutations were defined as uniquely observed in the evolved sample and not in the ancestral clone. Unassigned structural variants with good supporting evidence were added manually.

### Nucleotide diversity π of nonsynonymous and synonymous SNPs

To assess the selective pressures experienced by *P. putida* and *A. johnsonii* in monoculture versus co-culture environments, we calculated nucleotide diversity (π) for evolved populations from all detected single nucleotide polymorphisms (SNPs) at generation 800.

Since the ancestral state of our bacteria are known, nucleotide diversity can be calculated directly from derived allele frequencies to measure evolutionary divergence from the ancestor. For each polymorphic site detected by *breseq*, we used the derived allele frequency (*p*) as the measure of nucleotide diversity at that site (π = *p*), where *p* represents the frequency of the mutant allele relative to the known ancestral reference genome. This approach differs from classical population genetic measures of heterozygosity (π = 2*p*(1-*p*)) and instead quantifies the average degree of divergence from the ancestral state. Because the ancestral population was monomorphic at all sites, any polymorphism detected represents evolutionary change, and the frequency of derived alleles directly reflects the extent of divergence from the founder genotype.

To distinguish between different modes of selection, we separately calculated nucleotide diversity for nonsynonymous (π_N_) and synonymous (π_S_) mutations. Replicate experimental populations were pooled by treatment condition prior to calculating π_N_/π_S_ ratios, following approaches established for analyzing haploid microbial populations (Barrick and Lenski, 2009; Tenaillon et al., 2012).

The π_N_/π_S_ ratio serves as an indicator of selection regime: values less than 1 indicate that nonsynonymous mutations have accumulated to lower average frequencies than synonymous mutations, consistent with purifying selection removing deleterious mutations; values approximately equal to 1 suggest neutral evolution or relaxed selection; and values greater than 1 indicate that nonsynonymous mutations have reached higher average frequencies, potentially reflecting adaptive evolution or substantially relaxed purifying selection (Kryazhimskiy and Plotkin, 2008). In the context of experimental evolution, these ratios reflect the differential accumulation rates of nonsynonymous versus synonymous mutations over the course of the experiment.

### Statistical analysis of selection, enrichment and parallelism

We tested whether mutations that reached high frequency (>80%) by the endpoint of the evolution experiment were enriched by selection using the statistical methods previously described in (Rodríguez-Verdugo and Ackermann, 2021). Briefly, the strength of parallelism at the gene level across experimental lines within a condition was estimated using a Poisson distribution: with λ = total observed mutations genome size in bp × size of the target gene in bp. The p-value was estimated from the Poisson cumulative expectation, P (x ≥ observed, 1).

Gene Ontology (GO) enrichment analysis was performed to identify overrepresented biological functions within a gene set with fixed mutations within a condition across all experimental lines compared to the reference genome using ShinyGO (version 0.85). We used *P. putida* K2440 reference and the GO biological processes database with 0.05 FDR cutoff and redundancy removal. For *A. johnsonii*, we used a custom script that extracts the GO terms predicted in our assembled genome. GO terms were extracted directly from the GenBank annotation file using the Biopython package (version 1.81) in Python 3. For each gene in our list of mutated genes, associated GO terms were parsed from the feature note fields. Enrichment analysis was conducted using Fisher’s exact test (two-tailed) implemented in SciPy (version 1.11.0). For each GO term, a 2×2 contingency table was constructed comparing the number of genes with and without the term in the test set versus the reference genome. Fold enrichment was calculated as (*k*/*n*)/(*K*/*N*), where *k* represents genes in the test set with the GO term, *n* is the total number of genes in the gene set, *K* represents genes in the reference with the GO term, and *N* is the total number of genes in the reference genome. To correct for multiple hypothesis testing, p-values were adjusted using the Benjamini-Hochberg false discovery rate (FDR) method via the *statsmodels* package (version 0.14.0). GO terms with an adjusted p-value (q-value) < 0.05 were considered significantly enriched.

### Lineage tracking

To complement population-level data and resolve individual genotypes within lineages, we isolated eight clones per population per timepoint. Each unique combination of mutations from clones defined a distinct genotype. Mutations observed in the clones but not detected in population data were assumed to be rare and were not included in the lineage tracking. To link clonal genotypes to population lineages, we matched fixed mutations in clones to their frequency trajectories in population sequencing data. Mutations with correlated dynamics were assigned to the same lineage, with the earliest detected mutation serving as the lineage identifier. For each lineage, we calculated the total lineage frequency at each time point by summing the frequencies of all mutations within that lineage. Clonal data validated lineage assignments when multiple clones sharing identical mutation combinations corresponded to a single population lineage.

Muller plots visualizing lineage dynamics over evolutionary time were generated using the *ggmuller* package v0.5.6 in R v4.3.1. Input data consisted of lineage frequencies across all sampled time points. To construct proper parent-child relationships for nested lineages, we created adjacency tables defining the phylogenetic structure. Lineages arising from the ancestral background were designated as independent, while lineages sharing subsets of mutations indicated nested relationships. We manually curated the phylogeny based on mutation overlap: if lineage B contained all mutations from lineage A plus additional variants, lineage B was defined as a descendant of lineage A.

### Identification and analysis of large deletions in evolved *A. johnsonii*

The genomic region spanning coordinates 850,041-922,000 bp (71,960 bp) of the *A. johnsonii* strain C6 ancestral genome, encompassing the tmRNA gene (ssrA) and the deleted region observed in evolved populations, was extracted for detailed analysis. Genomic islands within this region were predicted using IslandViewer4 (Bertelli et al., 2017), which integrates three complementary prediction methods: IslandPath-DIMOB, SIGI-HMM, and IslandPick. The assembled C6 genome was aligned to *A. johnsonii* XBB1, the most closely related available reference genome.

To identify direct repeats flanking the deleted region that would indicate site-specific recombination, we performed pairwise sequence comparisons between the first and last 500 bp of the extracted sequence, searching for perfect matches of 8-50 bp in length using custom Python scripts. GC content was calculated across the region using a 1,000 bp sliding window with 100 bp steps. GC skew, defined as (G-C)/(G+C), was calculated using a 10,000 bp sliding window with 1,000 bp steps to detect potential replication strand bias or compositional heterogeneity. Statistical significance of GC composition differences between genomic islands and flanking regions was assessed using chi-square contingency tests.

The tmRNA (ssrA) gene was located by searching for conserved tmRNA sequence motifs. Prophage-associated sequences were identified by searching for integrase core binding sites and attachment (att) site motifs commonly found in prophage integration regions. All sequence analyses were performed using Python 3 with *NumPy* (Bertelli et al., 2017), *SciPy* (Bertelli et al., 2017), and *Matplotlib* (Hunter, 2007) libraries.

### Quantification of the evolution of ecological interactions

To measure possible changes in interactions between *P. putida* and *A. johnsonii*, we revived co-cultures Co1, Co2, and Co3 from frozen glycerol stocks at generations 6, 400, and 800. We derived monocultures of both species from these co-cultures by using selective antibiotics gentamicin and streptomycin to kill off the other species. The purity of the derived monocultures was confirmed through plating undiluted cultures on LB. Co-cultures revived from glycerol stocks were allowed to acclimate for 24 h in the same growth conditions as the evolution experiment. 100 µL were then inoculated into 10 mL of FAB with 0.6 mM of benzyl alcohol with streptomycin, gentamicin, or no antibiotics to create *A johnsonii* monoculture, *P. putida* monoculture, and a co-culture respectively. These cultures were grown for four days with 1% transfers of saturated media into fresh media every day to capture their growth dynamics after the removal of the other species. CFU counts were obtained at the end of each 24 h growth cycle. This experiment was conducted on three separate occasions to obtain a total of three temporal replicates

We calculated the area under the curve (AUC) using the trapezoidal method, integrating density over the 4-day period as a measure of overall growth. We performed paired t-tests comparing co-culture vs. monoculture AUC values for each generation × species × experimental line combination (n = 18 tests total), with temporal replicates paired by ID. P-values were adjusted using the Benjamini-Hochberg false discovery rate correction.

To test whether interaction strength evolved over time, we fit linear mixed-effects models with the difference in AUC (co-culture - monoculture) as the response variable, generation as a continuous fixed effect (6, 400, 800), and experimental line as a random effect. This was performed separately for each species.

## Data and resource availability

The genome sequence for *A. johnsonii* generated during the current study is available in GenBank (BioProject PRJNA1339041; BioSample SAMN52625548) under the accession numbers JBTYHZ010000001.1 and JBTYHZ010000002.1. All the other data underlying this article are available in the article and in its online supplementary material.

## Acknowledgments

We thank Brandon Gaut, Jennifer Martiny, and the Rodriguez-Verdugo lab members for providing useful feedback during this research. This work was supported by the National Science Foundation (Grant no. DEB-2234627). Z.A. was supported by the NSF GAANN fellowship.

## Notes

### Competing Interest Statement

The authors have declared no competing interest.

