## Supplementary Materials for "Unilateral cross-feeding constrains adaptive evolution, even in the producer without direct fitness effects"

### Supplementary Figures

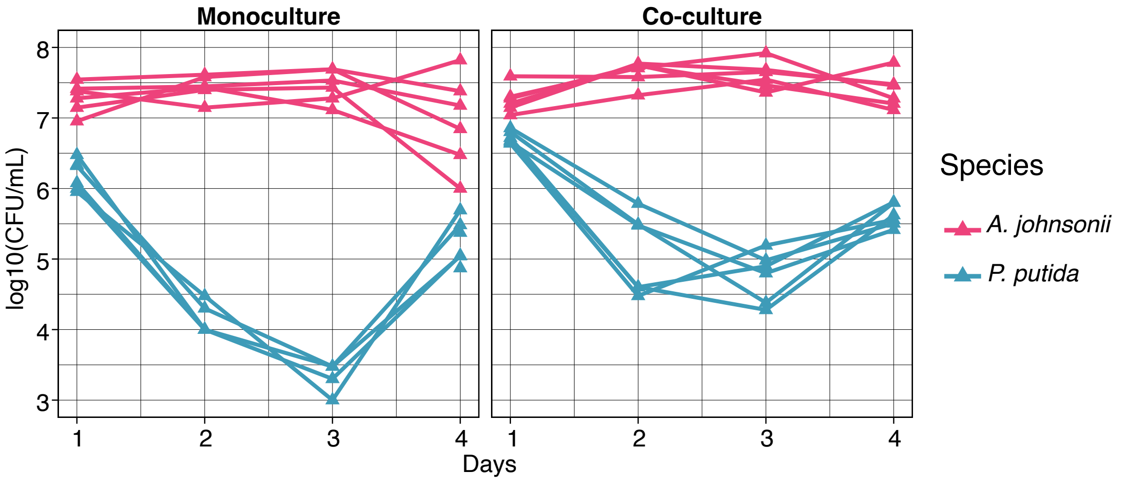


**Figure S1.** Population density (CFU mL$^{-1}$) trajectories of six replicates of *A. johnsonii* and *P. putida* ancestors grown in monoculture (left panel) and co-culture (right panel) over four days of serial transfers.

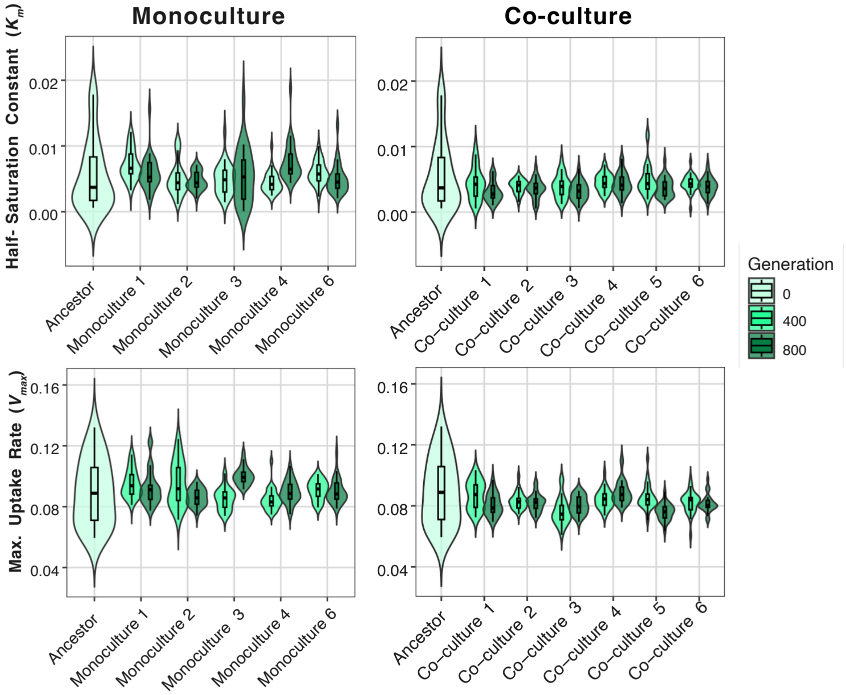


**Figure S2.** Half-saturation constant ($K_m$) and maximum uptake rate ($V_{max}$) for monoculture (left panel) and co-culture (right panel) evolved *P. putida*.

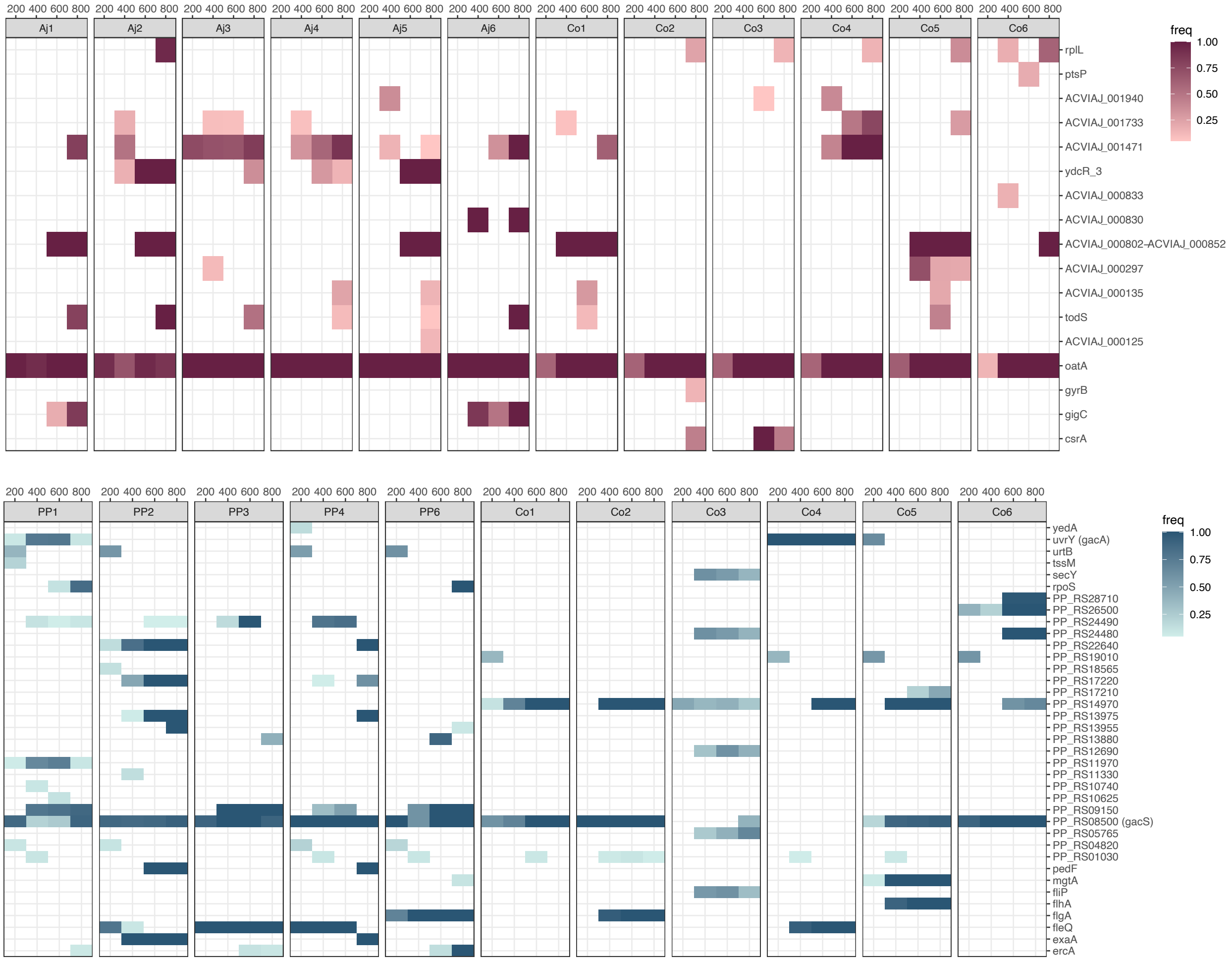


**Figure S3.** Heat map showing the frequency of nonsynonymous mutations (genes = rows) across generations in populations of *A. johnsonii* (red) and *P. putida* (blue). Six replicate monocultures (Aj1-Aj6 and PP1-PP6) and co-cultures (Co1-Co6) are shown. Only genes with mutations at frequencies above 0.1 are shown. The frequency of each mutation in the population is shown by different intensities of blue.

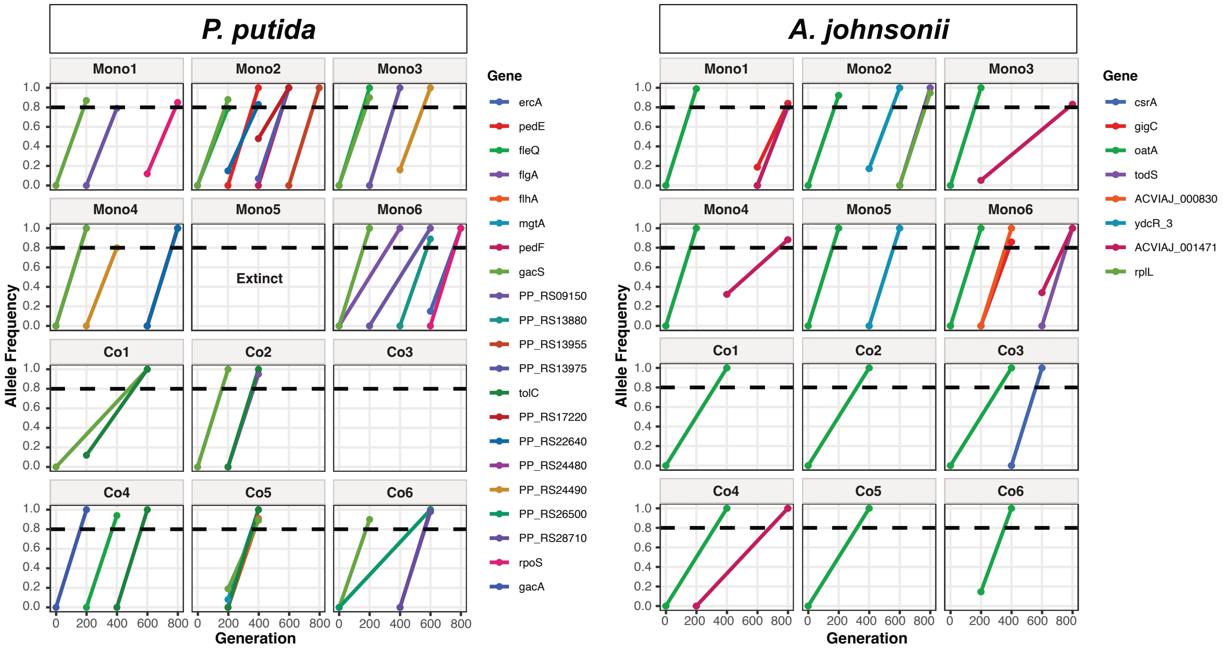


**Figure S4.** Selective sweeps of mutations reaching over 80% frequency in populations of *P. putida* (left panel) and *A. johnsonii* (right panel). Six replicate monocultures (Mono1-Mono6) and co-cultures (Co1-Co6) are shown.

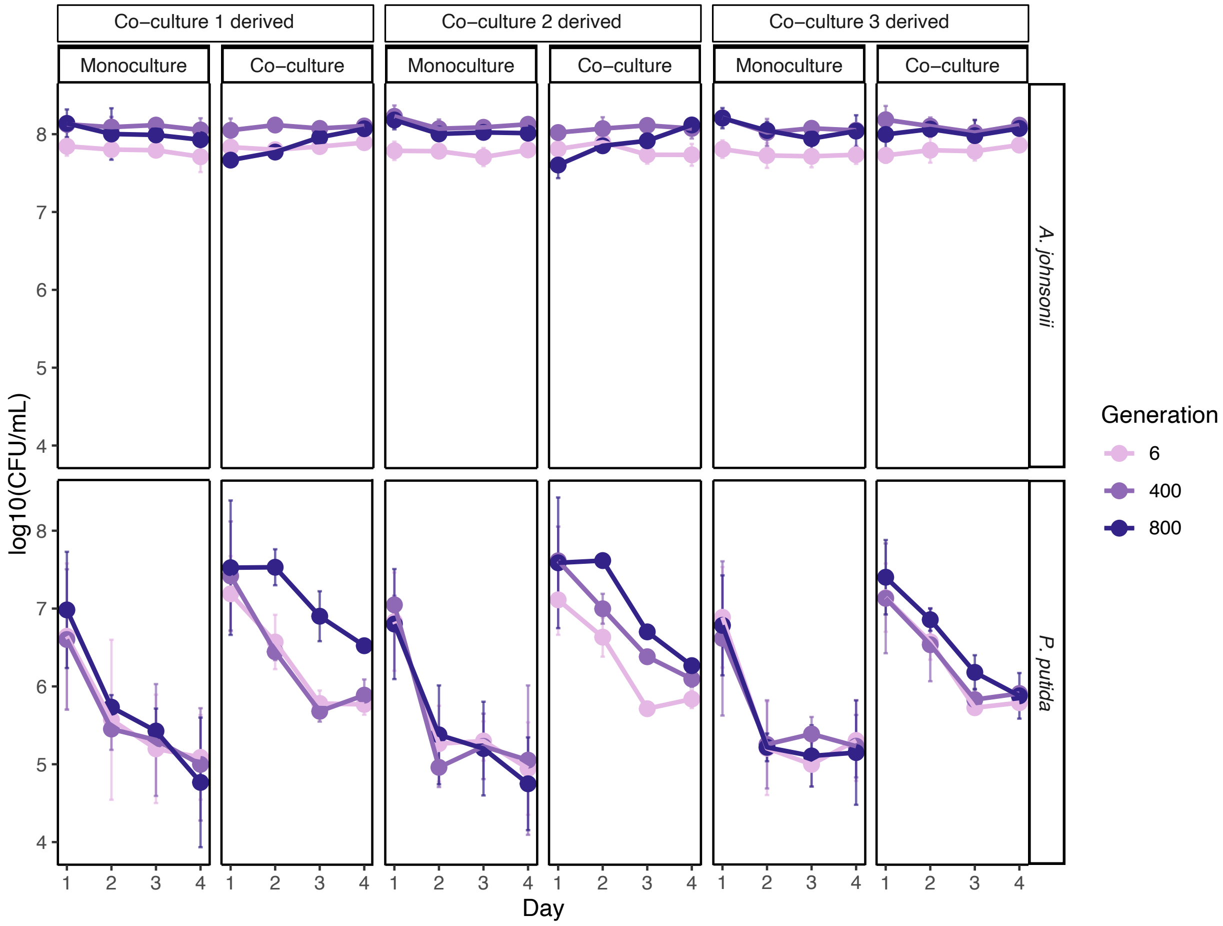


**Figure S5.** Growth dynamics of three co-cultures revived from generation 6, 400, and 800 timepoints. Monocultures were derived from co-cultures (Co1 to Co3) using antibiotics. The area under the curve (AUC) was used to assess the growth of each species under each condition and across generations (Table S7). *P. putida* exhibited strong and consistent co-culture benefits across all generations and experimental lines (9/9 comparisons significant after FDR correction, $p < 0.024$ for all). Mean co-culture advantage ranged from +2.19 AUC units at 6 generations to +4.89 at 800 generations, representing substantial growth enhancement. *A. johnsonii* showed no significant co-culture effect at any generation (0/9 comparisons significant after FDR correction, $p > 0.17$ for all). Mean differences were near zero or slightly negative across all time points.

### Supplementary Tables

**Table S1.** Resource use traits fold-change differences between monocultures and co-cultures for *Pseudomonas putida*. Values represent fold-change relative to the ancestor. Statistical comparisons by Welch's *t*-test.

| Trait | Generation | $n_{mono}$ | $n_{co}$ | $Mean_{mono}$ | $SE_{mono}$ | $Mean_{co}$ | $SE_{co}$ | Difference | $t$ | $df$ | $P$ | Cohen's $d$ | |
|---|---|---|---|---|---|---|---|---|---|---|---|---|---|
| Yield | 400 | 5 | 6 | 2.882 | 0.071 | 2.221 | 0.262 | 0.661 | 2.431 | 9 | 0.053 | 1.347 | — |
| Yield | 800 | 5 | 6 | 3.186 | 0.145 | 2.469 | 0.131 | 0.717 | 3.674 | 9 | 0.006 | 2.228 | ** |
| $\mu_{max}$ | 400 | 5 | 6 | 2.834 | 0.037 | 2.141 | 0.271 | 0.693 | 2.536 | 9 | 0.050 | 1.393 | — |
| $\mu_{max}$ | 800 | 5 | 6 | 3.037 | 0.122 | 2.336 | 0.147 | 0.701 | 3.677 | 9 | 0.005 | 2.166 | ** |
| $K$ | 400 | 5 | 6 | 0.973 | 0.084 | 0.771 | 0.027 | 0.203 | 2.288 | 9 | 0.073 | 1.500 | — |
| $K$ | 800 | 5 | 6 | 1.018 | 0.091 | 0.648 | 0.034 | 0.370 | 3.823 | 9 | 0.012 | 2.489 | * |
| $V_{max}$ | 400 | 5 | 6 | 1.009 | 0.026 | 0.930 | 0.017 | 0.079 | 2.567 | 9 | 0.036 | 1.603 | * |
| $V_{max}$ | 800 | 5 | 6 | 1.026 | 0.026 | 0.913 | 0.019 | 0.113 | 3.502 | 9 | 0.009 | 2.173 | ** |

*Note:* $\mu_{max}$ = maximum growth rate; $K$ = half-saturation constant; $V_{max}$ = maximum uptake rate. Mono = monoculture; Co = coculture.
* $P < 0.05$; ** $P < 0.01$; — not significant.

**Table S2.** Pairwise comparisons of resource use traits across generations. A separate file (.xlsx) is provided.

**Table S3.** *de novo* mutations in populations and clones identified using *breseq*. A separate file (.xlsx) is provided.

**Table S4.** Nucleotide diversity ($\pi$) summary for nonsynonymous and synonymous sites in evolved populations of *Pseudomonas putida* and *Acinetobacter johnsonii* under monoculture and coculture conditions.

| Species | Condition | $n$ $sites_N$ | $n$ $sites_S$ | $\Sigma\pi_N$ | $\Sigma\pi_S$ | Mean $\pi_N$ | Mean $\pi_S$ | $\pi_N/\pi_S$ |
|---|---|---|---|---|---|---|---|---|
| *A. johnsonii* | Co-culture | 105 | 149 | 9.096 | 12.364 | 0.0866 | 0.0830 | 1.044 |
| | Monoculture | 99 | 61 | 14.939 | 4.552 | 0.1509 | 0.0746 | 2.022 |
| *P. putida* | Co-culture | 11 | 10 | 5.490 | 6.620 | 0.4991 | 0.6620 | 0.754 |
| | Monoculture | 343 | 230 | 29.610 | 13.820 | 0.0863 | 0.0601 | 1.437 |

*Note:* $\pi_N$ = nucleotide diversity at nonsynonymous sites; $\pi_S$ = nucleotide diversity at synonymous sites; $\Sigma$ = cumulative sum across all mutated genes; $\pi_N/\pi_S > 1$ indicates an excess of nonsynonymous diversity consistent with positive selection while $\pi_N/\pi_S < 1$ indicates purifying selection.

**Table S5.** Significantly enriched Gene Ontology (GO) pathways among genes harboring mutations in evolved populations of *Pseudomonas putida* and *Acinetobacter johnsonii*. Enrichment was assessed by Fisher's exact test with FDR correction (Benjamini–Hochberg).

| Condition | Species | GO pathway | $\log_2$(fold enrichment) | FDR |
|---|---|---|---|---|
| Coculture | *P. putida* | Biological regulation | 2.149 | $3.90 \times 10^{-2}$ |
| | | Peptidyl-amino acid modification | 4.540 | $3.62 \times 10^{-2}$ |
| | | Peptidyl-histidine modification | 5.083 | $2.39 \times 10^{-2}$ |
| | | Peptidyl-histidine phosphorylation | 5.083 | $2.39 \times 10^{-2}$ |
| | | Protein metabolic process | 2.925 | $3.90 \times 10^{-2}$ |
| | | Protein modification process | 3.911 | $4.46 \times 10^{-2}$ |
| | | Protein phosphorylation | 4.816 | $2.96 \times 10^{-2}$ |
| | | Regulation of biological process | 2.257 | $3.62 \times 10^{-2}$ |
| | | Regulation of cell motility | 6.442 | $6.36 \times 10^{-3}$ |
| | | Regulation of locomotion | 6.322 | $6.36 \times 10^{-3}$ |
| Monoculture | *P. putida* | Bacterial-type flagellum assembly | 6.480 | $5.45 \times 10^{-3}$ |
| | | Bacterial-type flagellum organization | 5.764 | $7.42 \times 10^{-3}$ |
| | | Cell projection assembly | 5.334 | $1.06 \times 10^{-2}$ |
| | | Cell projection organization | 4.930 | $1.72 \times 10^{-2}$ |
| | | Cellular component assembly | 3.560 | $4.18 \times 10^{-2}$ |
| | | Non-membrane-bounded organelle assembly | 5.764 | $7.42 \times 10^{-3}$ |
| | | Nucleic acid-templated transcription | 4.288 | $3.19 \times 10^{-2}$ |
| | | Organelle assembly | 5.764 | $7.42 \times 10^{-3}$ |
| | | Organelle organization | 4.002 | $3.19 \times 10^{-2}$ |
| | | Regulation of cell adhesion | 7.288 | $3.19 \times 10^{-2}$ |
| | | Regulation of cell motility | 6.349 | $2.74 \times 10^{-4}$ |
| | | Regulation of cell-substrate adhesion | 7.703 | $3.19 \times 10^{-2}$ |
| | | Regulation of cellular component biogenesis | 6.966 | $3.71 \times 10^{-2}$ |
| | | Regulation of cellular response to stress | 6.480 | $4.18 \times 10^{-2}$ |
| | | Regulation of chemotaxis | 7.288 | $3.19 \times 10^{-2}$ |
| | | Regulation of locomotion | 6.229 | $2.74 \times 10^{-4}$ |
| | | Regulation of response to external stimulus | 6.480 | $4.18 \times 10^{-2}$ |
| | | Regulation of response to oxidative stress | 7.288 | $3.19 \times 10^{-2}$ |
| | | Regulation of secondary metabolic process | 5.644 | $7.42 \times 10^{-3}$ |
| | | Regulation of secondary metabolite biosynthetic process | 5.644 | $7.42 \times 10^{-3}$ |
| | | RNA biosynthetic process | 4.138 | $3.19 \times 10^{-2}$ |
| | | Transcription, DNA-templated | 4.288 | $3.19 \times 10^{-2}$ |
| Coculture | *A. johnsonii* | Acyltransferase activity, transferring groups other than amino-acyl groups | 7.040 | $7.58 \times 10^{-3}$ |
| Monoculture | *A. johnsonii* | Acyltransferase activity, transferring groups other than amino-acyl groups | 5.455 | $4.52 \times 10^{-2}$ |

*Note:* $\log_2$(fold enrichment) indicates the magnitude of overrepresentation relative to the genomic background. Only pathways with FDR < 0.05 are shown. Rows are sorted alphabetically within each condition × species group.

**Table S6.** Predicted genomic islands in *A. johnsonii* C6 identified by IslandViewer 4. Gene content for each island is listed with locus tags, coordinates, and predicted products.

| Island | Coordinates | Length (bp) | Prediction method | Locus tag | Start | End | Strand | Product |
|---|---|---|---|---|---|---|---|---|
| 1 | 856,233–861,527 | 5,294 | Predicted by ≥1 method | ACVIAJ_000809 | 856,233 | 856,508 | − | hypothetical protein |
| | | | | ACVIAJ_000810 | 857,335 | 857,652 | + | hypothetical protein |
| | | | | ACVIAJ_000811 | 857,691 | 858,737 | + | DUF932 domain-containing protein |
| | | | | ACVIAJ_000812 | 858,887 | 859,930 | + | YqaJ viral recombinase family protein |
| | | | | ACVIAJ_000813 | 860,048 | 860,977 | + | hydrolase or metal-binding protein |
| | | | | ACVIAJ_000814 | 861,045 | 861,275 | + | helix-turn-helix transcriptional regulator |
| | | | | ACVIAJ_000815 | 861,348 | 861,527 | + | hypothetical protein |
| 2 | 889,088–900,395 | 11,307 | Predicted by ≥1 method | ACVIAJ_000830 | 889,088 | 889,465 | + | helix-turn-helix domain-containing protein |
| | | | | ACVIAJ_000831 | 889,472 | 890,296 | − | hypothetical protein |
| | | | | ACVIAJ_000832 | 891,276 | 892,127 | + | OmpA family protein |
| | | | | ACVIAJ_000833 | 892,170 | 900,395 | + | BapA/Bap/LapF family prefix-like domain-containing protein |
| 3 | 908,174–921,589 | 13,415 | Predicted by ≥1 method | ACVIAJ_000840 | 908,174 | 908,473 | + | recombinase-like helix-turn-helix domain-containing protein |
| | | | | ACVIAJ_000841 | 908,489 | 909,553 | + | Rieske 2Fe-2S domain-containing protein |
| | | | | ACVIAJ_000842 | 909,546 | 910,310 | + | SDR family oxidoreductase |
| | | | | ACVIAJ_000843 | 910,321 | 911,250 | + | VOC family protein |
| | | | | ACVIAJ_000844 | 911,276 | 912,484 | + | NAD(P)/FAD-dependent oxidoreductase |
| | | | | ACVIAJ_000845 | 912,525 | 913,748 | + | MFS transporter |
| | | | | ACVIAJ_000846 | 913,914 | 914,435 | + | cupin domain-containing protein |
| | | | | ACVIAJ_000847 | 914,451 | 915,302 | + | alpha/beta fold hydrolase |
| | | | | ACVIAJ_000848 | 915,312 | 916,106 | + | SDR family oxidoreductase |
| | | | | ACVIAJ_000849 | 916,116 | 916,907 | + | aspartate dehydrogenase |
| | | | | ACVIAJ_000850 | 916,927 | 918,393 | + | aldehyde dehydrogenase |
| | | | | ACVIAJ_000851 | 918,442 | 920,082 | + | thiamine pyrophosphate-binding protein |
| | | | | ACVIAJ_000852 | 920,390 | 921,589 | + | hypothetical protein |

*Note:* Islands were predicted by at least one of the integrated methods in IslandViewer 4 (IslandPath-DIMOB, SIGI-HMM, or IslandPick). Coordinates refer to positions in the complete chromosome (GenBank accession PRJNA1339041). Strand: +, sense; −, antisense.

**Table S7.** Calculated area under the curve (AUC)

| Generation | Species | Experimental line | Replicate | $AUC_{Co}$ | $AUC_{Mono}$ | Partner's effect ($AUC_{Co}$ - $AUC_{Mono}$) |
|---|---|---|---|---|---|---|
| 6 | *A. johnsonii* | co1 | 1 | 23.635 | 23.609 | 0.027 |
| 6 | *A. johnsonii* | co1 | 2 | 23.589 | 23.151 | 0.438 |
| 6 | *A. johnsonii* | co1 | 3 | 23.310 | 23.385 | -0.075 |
| 6 | *A. johnsonii* | co2 | 1 | 23.642 | 23.259 | 0.383 |
| 6 | *A. johnsonii* | co2 | 2 | 23.410 | 23.347 | 0.063 |
| 6 | *A. johnsonii* | co2 | 3 | 23.186 | 23.257 | -0.071 |
| 6 | *A. johnsonii* | co3 | 1 | 23.506 | 22.988 | 0.518 |
| 6 | *A. johnsonii* | co3 | 2 | 23.584 | 23.402 | 0.182 |
| 6 | *A. johnsonii* | co3 | 3 | 23.041 | 23.250 | -0.209 |
| 6 | *P. putida* | co1 | 1 | 18.847 | 17.391 | 1.456 |
| 6 | *P. putida* | co1 | 2 | 18.965 | 16.701 | 2.264 |
| 6 | *P. putida* | co1 | 3 | 18.705 | 15.840 | 2.865 |
| 6 | *P. putida* | co2 | 1 | 18.900 | 16.634 | 2.266 |
| 6 | *P. putida* | co2 | 2 | 18.786 | 16.350 | 2.436 |
| 6 | *P. putida* | co2 | 3 | 18.804 | 16.396 | 2.408 |
| 6 | *P. putida* | co3 | 1 | 18.739 | 16.652 | 2.087 |
| 6 | *P. putida* | co3 | 2 | 18.782 | 15.994 | 2.788 |
| 6 | *P. putida* | co3 | 3 | 18.801 | 16.253 | 2.548 |
| 400 | *A. johnsonii* | co1 | 1 | 24.397 | 24.481 | -0.084 |
| 400 | *A. johnsonii* | co1 | 2 | 24.377 | 24.265 | 0.111 |
| 400 | *A. johnsonii* | co1 | 3 | 24.059 | 24.172 | -0.114 |
| 400 | *A. johnsonii* | co2 | 1 | 24.329 | 24.330 | 0.000 |
| 400 | *A. johnsonii* | co2 | 2 | 24.310 | 24.413 | -0.103 |
| 400 | *A. johnsonii* | co2 | 3 | 24.080 | 24.292 | -0.212 |
| 400 | *A. johnsonii* | co3 | 1 | 24.610 | 24.046 | 0.565 |
| 400 | *A. johnsonii* | co3 | 2 | 24.319 | 24.393 | -0.074 |
| 400 | *A. johnsonii* | co3 | 3 | 23.918 | 24.244 | -0.326 |
| 400 | *P. putida* | co1 | 1 | 18.433 | 16.631 | 1.802 |
| 400 | *P. putida* | co1 | 2 | 18.932 | 16.752 | 2.180 |
| 400 | *P. putida* | co1 | 3 | 18.998 | 16.336 | 2.662 |
| 400 | *P. putida* | co2 | 1 | 20.196 | NA | NA |
| 400 | *P. putida* | co2 | 2 | 20.396 | 16.495 | 3.902 |
| 400 | *P. putida* | co2 | 3 | 20.134 | 16.327 | 3.807 |
| 400 | *P. putida* | co3 | 1 | 19.017 | 16.651 | 2.366 |
| 400 | *P. putida* | co3 | 2 | 18.946 | 16.664 | 2.282 |
| 400 | *P. putida* | co3 | 3 | 18.724 | 16.393 | 2.332 |
| 800 | *A. johnsonii* | co1 | 1 | 23.653 | 24.377 | -0.724 |
| 800 | *A. johnsonii* | co1 | 2 | 23.647 | 24.045 | -0.397 |
| 800 | *A. johnsonii* | co1 | 3 | 23.518 | 23.682 | -0.164 |
| 800 | *A. johnsonii* | co2 | 1 | 23.468 | 24.241 | -0.773 |
| 800 | *A. johnsonii* | co2 | 2 | 23.896 | 24.102 | -0.206 |
| 800 | *A. johnsonii* | co2 | 3 | 23.541 | 24.053 | -0.511 |
| 800 | *A. johnsonii* | co3 | 1 | 24.369 | 24.257 | 0.112 |
| 800 | *A. johnsonii* | co3 | 2 | 24.282 | 24.028 | 0.255 |
| 800 | *A. johnsonii* | co3 | 3 | 23.618 | 24.052 | -0.434 |
| 800 | *P. putida* | co1 | 1 | 20.648 | 16.387 | 4.261 |
| 800 | *P. putida* | co1 | 2 | 22.321 | 17.076 | 5.245 |
| 800 | *P. putida* | co1 | 3 | 21.426 | 17.667 | 3.759 |
| 800 | *P. putida* | co2 | 1 | 20.834 | 16.230 | 4.604 |
| 800 | *P. putida* | co2 | 2 | 21.457 | 16.856 | 4.601 |
| 800 | *P. putida* | co2 | 3 | 21.469 | 16.004 | 5.466 |
| 800 | *P. putida* | co3 | 1 | 19.103 | 15.778 | 3.325 |
| 800 | *P. putida* | co3 | 2 | 19.966 | 16.791 | 3.175 |
| 800 | *P. putida* | co3 | 3 | 19.993 | 16.304 | 3.688 |

**Table S8.** Linear mixed-effects model results for the effect of generation on the difference in area under the curve (AUC) between coculture and monoculture conditions. Separate models were fit for *A. johnsonii* and *P. putida*. Values in parentheses are standard errors.

| | ***A. johnsonii*** | ***P. putida*** |
|---|---|---|
| **Model specification** | | |
| Formula | *Difference* ~ *Generation* + (1\|*Line*) | *Difference* ~ *Generation* + (1\|*Line*) |
| Observations | 27 | 26 |
| Experimental lines | 3 | 3 |
| **Fixed effects** | | |
| Intercept | 0.160 (0.095) | 2.172 (0.324) |
| Intercept *P* | 0.135 | 0.006 ** |
| Generation slope | −0.00057 (0.00016) | +0.00236 (0.00037) |
| Generation *P* | 0.002 ** | < 0.001 *** |
| **Random effects ($\sigma^2$)** | | |
| Line (intercept) | 0.072 | 0.449 |
| Residual | 0.279 | 0.633 |
| **Interpretation** | | |
| Effect direction | Coculture benefit decreases over time | Coculture benefit increases over time |
| Rate of change | −0.00057 per generation | +0.00236 per generation |

*Note:* Experimental line was included as a random intercept. * $P < 0.05$; ** $P < 0.01$; *** $P < 0.001$.
